# Role of the C-terminal tail in regulating Proteinase Activated Receptor 2 (PAR2) signalling

**DOI:** 10.1101/2020.03.02.973842

**Authors:** Pierre E. Thibeault, Rithwik Ramachandran

**Author notes:** To whom correspondence should be addressed: Rithwik Ramachandran, Department of Physiology and Pharmacology, University of Western Ontario. London, ON, N6A 5C1.

## Abstract

The C-terminal tail of G-protein-coupled receptors contain important regulatory sites that enable interaction with intracellular signalling effectors. Here we examine the relative contribution of the C-tail serine/threonine phosphorylation sites (Ser^383-385^, Ser^387^-Thr^392^) and the helix-8 palmitoylation site (Cys^361^) in signalling regulation downstream of the proteolytically-activated GPCR, PAR2. We examined Gα_q/11_-coupled calcium signalling, β-arrestin-1/-2 recruitment, and MAPK activation (p44/42 phosphorylation) by wild-type and mutant receptors expressed in a CRISPR/Cas9 PAR2-knockout HEK-293 cell background. We find that alanine substitution of the membrane proximal serine residues (Ser^383-385^Ala) had no effect on SLIGRL-NH_2_- or trypsin-stimulated β-arrestin recruitment. Alanine substitutions in the Ser^387^-Thr^392^ cluster resulted in a large (∼50%) decrease in β-arrestin-1/2 recruitment triggered by the activating peptide, SLIGRL-NH_2,_ but was without effect on trypsin-activated β-arrestin-1/-2 recruitment. Additionally, we find that alanine substitution of the helix-8 cysteine residue (Cys^361^Ala) led to a (∼50%) decrease in β-arrestin-1/-2 recruitment in response to both trypsin and SLIGRL-NH_2_. We further show that Gα_q/11_inhibition with YM254890, inhibited ERK phosphorylation by PAR2 agonists, while genetic deletion of β-arrestin-1/-2 by CRISPR/Cas9 enhanced MAPK activation. Knockout of β-arrestins also enhanced Gα_q/11_-mediated calcium signalling. In line with these findings, C-tail serine/threonine and cysteine residue mutants that have decreased β-arrestin recruitment also showed enhanced ERK activation. Thus, our studies point to multiple mechanisms that regulate β-arrestin interaction with PAR2 to regulate receptor-mediated signalling.

---

G-protein-coupled receptors (GPCRs) are important cell surface receptors that regulate diverse physiological processes in response to a variety of extracellular stimuli (1). The proteinase activated receptors (PARs) are a four-member family of GPCRs that are activated by a wide range of proteolytic enzymes including coagulation cascade-, immune cell-, and pathogen-derived proteinases (2, 3). PAR activation occurs when proteolytic enzymes remove a portion of the receptor N-terminus to reveal a cryptic tethered-ligand which can bind intramolecularly to activate the receptor (3). Synthetic hexapeptide mimetics of the tethered ligand can also be used to selectively activate individual members of this family (4–7).

PAR2 is a trypsin-like, serine proteinase-activated, member of this family and is well established as a major regulator of inflammatory responses (8, 9). PAR2-targeted compounds are highly sought after as novel anti-inflammatory agents. In recent studies, there is also an emerging realization that different proteolytic enzymes cleave and activate PAR2 in a manner that favours differential coupling to various intracellular effectors (10). This has given impetus to studies investigating molecular mechanisms underlying biased signalling through this receptor.

In keeping with observations in a variety of other GPCRs, PAR2 can signal through both G-protein coupling or via interactions with the multi-functional scaffold proteins, β-arrestin-1 and −2 (11). PAR2 activation by the enzymatically (e.g. trypsin) revealed tethered-ligand or synthetic activating peptides (e.g SLIGRL-NH_2_) trigger robust Gα_q/11_ coupling; and activation of Gα_12/13_ by this receptor has also been reported (12). Some non-canonical enzymatic activators of PAR2 are also reported to cause receptor coupling to other G-proteins (13). Additionally, PAR2 is a strong recruiter of β-arrestin-1/-2 in response to both tethered-ligand and peptide activation (14, 15).

β-arrestin-1/-2 interaction with activated GPCRs has a role in desensitizing the G-protein-mediated signalling event. Moreover, β-arrestins can also act as scaffolds for other intracellular signalling effectors that are activated by GPCRs (16) including PAR2 (11). β-arrestin interaction with activated GPCRs is believed to occur in a two-step manner with the first contacts at GPCR kinase (GRKs) phosphorylated serine and threonine residues in the C-terminal tail of the receptors (17, 18) which is followed by additional stable interactions with the receptor transmembrane core (19, 20).

Previous studies have examined the consequence of modifying the C-terminal tail of PAR2. Deletion of certain segments in the PAR2 C-tail (Ala^355^-Ser^363^; three-letter amino acid codes used throughout) or truncation of the tail (His^354^-Stop) resulted in a loss of agonist-mediated internalization and inositol 1,4,5-trisphosphate (IP_3_) accumulation (21). In the same study deletion of the PAR2 C-tail Ala^355^-Ser^363^ segment did not however affect ERK activation, while His^354^-Stop mutation did have an effect on this signalling pathway (21). The truncation mutations described above encompass two key regulatory sites, clusters of serine/threonine residues that are sites of phosphorylation and a cysteine residue that is a site for palmitoylation.

Mutational studies expressing PAR2 with various alanine substitutions to clusters of serine/threonine residues in Rat1 fibroblasts and HeLa cells have clearly established the C-tail as the site for phosphorylation and demonstrated that these sites are key for β-arrestin-mediated receptor desensitization (22).

The role of the C-terminal cysteine residue that is part of the truncations described above have also been probed and Cys^361^ was established as the primary site of palmitoylation in PAR2 (23, 24). In these studies, CHO cells expressing a Cys^361^ mutant curiously showed decreased calcium signalling but enhanced ERK activation in response to both trypsin and the activating peptide SLIGRL-NH_2_ (23); and also showed reduction in agonist-induced β-arrestin recruitment, receptor endocytosis and degradation (24).

Together, these studies have pointed to multiple regulatory mechanisms that are dependent on PAR2 C-tail that can regulate receptor-mediated signalling and desensitization. These studies have, however, been conducted in different cell lines and in some cases have utilized cells that endogenously express low levels of PAR2. Here we attempt to gain a more direct comparison of the contribution of C-terminal tail sites to PAR2 signalling. We used CRISPR/Cas9 targeting to generate a functional PAR2-knockout HEK-293 cell line and studied signalling responses to reconstituted wt-PAR2 and four mutated PAR2 receptors with alanine substitutions in C-terminal serine/threonine or cysteine residues. We find that several such modifications have a significant impact on β-arrestin recruitment to PAR2. Additionally, PAR2-mediated calcium signalling is desensitized less efficiently in cells that express mutants that are not able to recruit β-arrestin efficiently. In agreement with this finding, PAR2 desensitization is delayed in β-arrestin-1/-2 deficient HEK cells. We also find that MAPK activation (p44/42 phosphorylation; ERK activation) by PAR2 is dependent on Gα_q/11_-mediated signalling and was exaggerated in cells expressing mutated receptors that did not recruit β-arrestin-1/-2 comparably. These studies further clarify mechanisms underlying PAR2-dependent signalling.

## RESULTS

### SLIGRL-NH_2_-stimulated β-arrestin recruitment is decreased in Cys^361^Ala, Ser^387^-Thr^392^Ala, and Ser^383-385^Ala/Ser^387^-Thr^392^Ala mutants, but is unaffected in the Ser^383-385^Ala mutants

To investigate the role of C-tail mutations on the recruitment of β-arrestins, we employed bioluminescence resonance energy transfer (BRET) using C-terminally fused receptor/enhanced yellow fluorescent protein constructs as our photon acceptor and β-arresin-1-rluc (β-arrestin-1) or β-arresin-2-rluc (β-arrestin-2) as the photon donor. We determined the EC_50_ for SLIGRL-NH_2_-stimulated recruitment of β-arrestin-1 and −2 to wildtype PAR2-YFP as 12.1 ± 1.6 µM and 11.1 ± 1.2 µM, respectively (Fig. 2). We first assessed recruitment of β-arrestins to PAR2 with Cys^361^Ala mutation (PAR2^C361A^-YFP) to determine the impact of loss of a known palmitoylation site in the C-tail of PAR2. We observed that SLIGRL-NH_2_-stimulated recruitment of both β-arrestin-1 and −2 recruitment to PAR2^C361A^-YFP are significantly decreased compared to PAR2-YFP (EC_50_ = 28.8 ± 4.0 µM and 27.4 ± 2.5 µM, respectively). Additionally, we observe that there is a significant decrease in maximal net BRET (Max.; top of the curve utilized for nonlinear regression curve fitting) of PAR2^C361A^-YFP (β-arrestin-1 Max. = 0.28 ± 0.01; β-arrestin-2 Max. = 0.32 ± 0.01) compared to PAR2-YFP (β-arrestin-1 Max. = 0.39 ± 0.01; β-arrestin-2 Max. = 0.53 ± 0.01) (Fig. 2).

**Figure 1:**
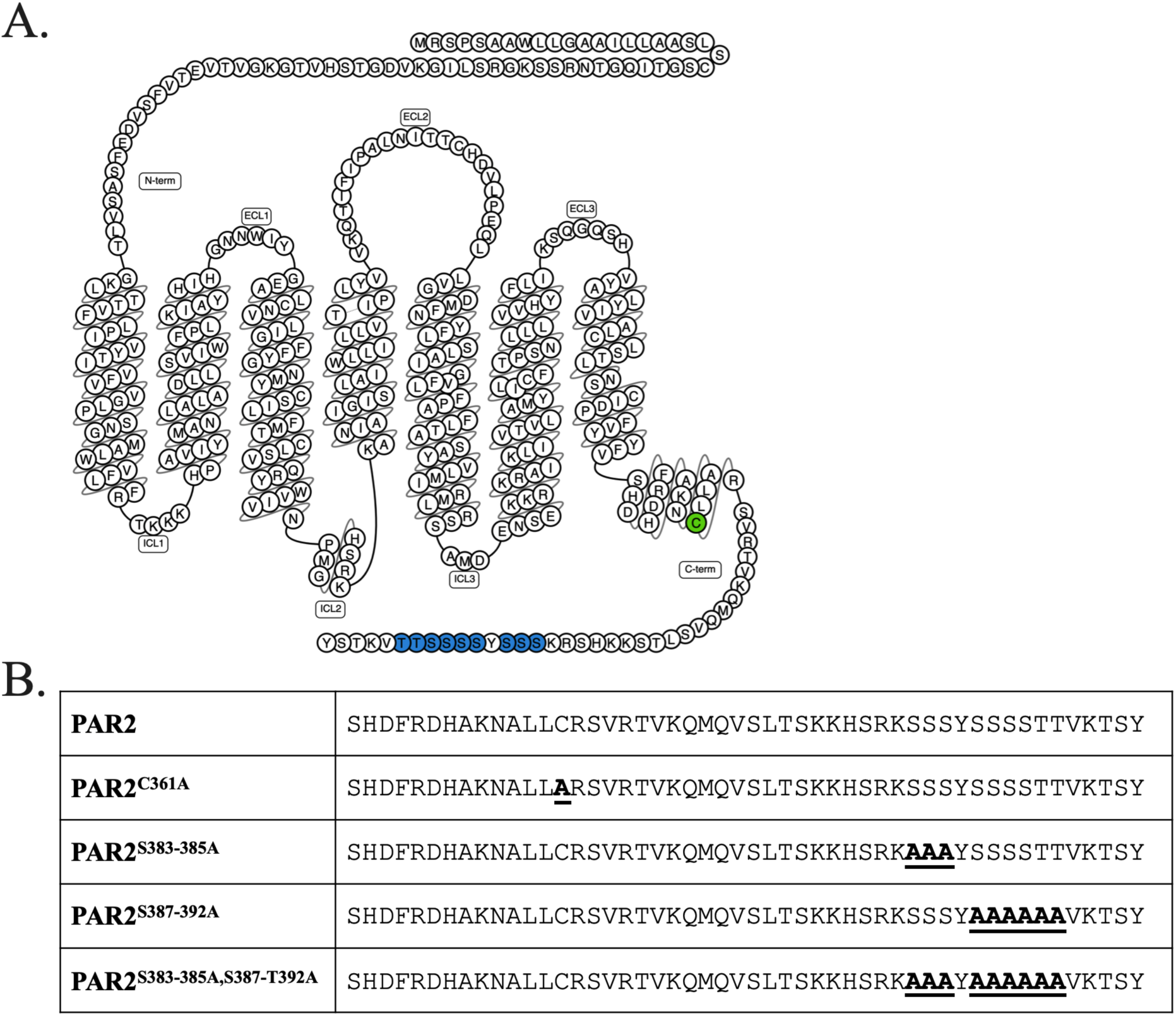
Summary of PAR2 C-tail residue mutations. (A) Cartoon depicting PAR2 with palmitoylation site (green) and phosphorylation sites (blue) shown (generated with GPCRdb) (67, 68). (B) Mutant receptor clones generated using Agilent QuikChange XL site-directed mutagenesis. Sequences of the wildtype PAR2 and mutant PAR2 receptor C-tail constructs (mutations are shown underlined). PAR2C^361^A mutation was generated to study the effects of this residue on signalling endpoints. PAR2S^383-386^A and PAR2S^387-^T^392A^ mutants were generated to determine the effects of partial mutation of the serine/threonine rich region of the PAR2 C-tail. A combined mutant of PAR2S^383-385^A, S^387-^T^392^A was also generated to determine the effect on mutation of serine/threonine rich C-tail motif on signalling.

**Figure 2:**
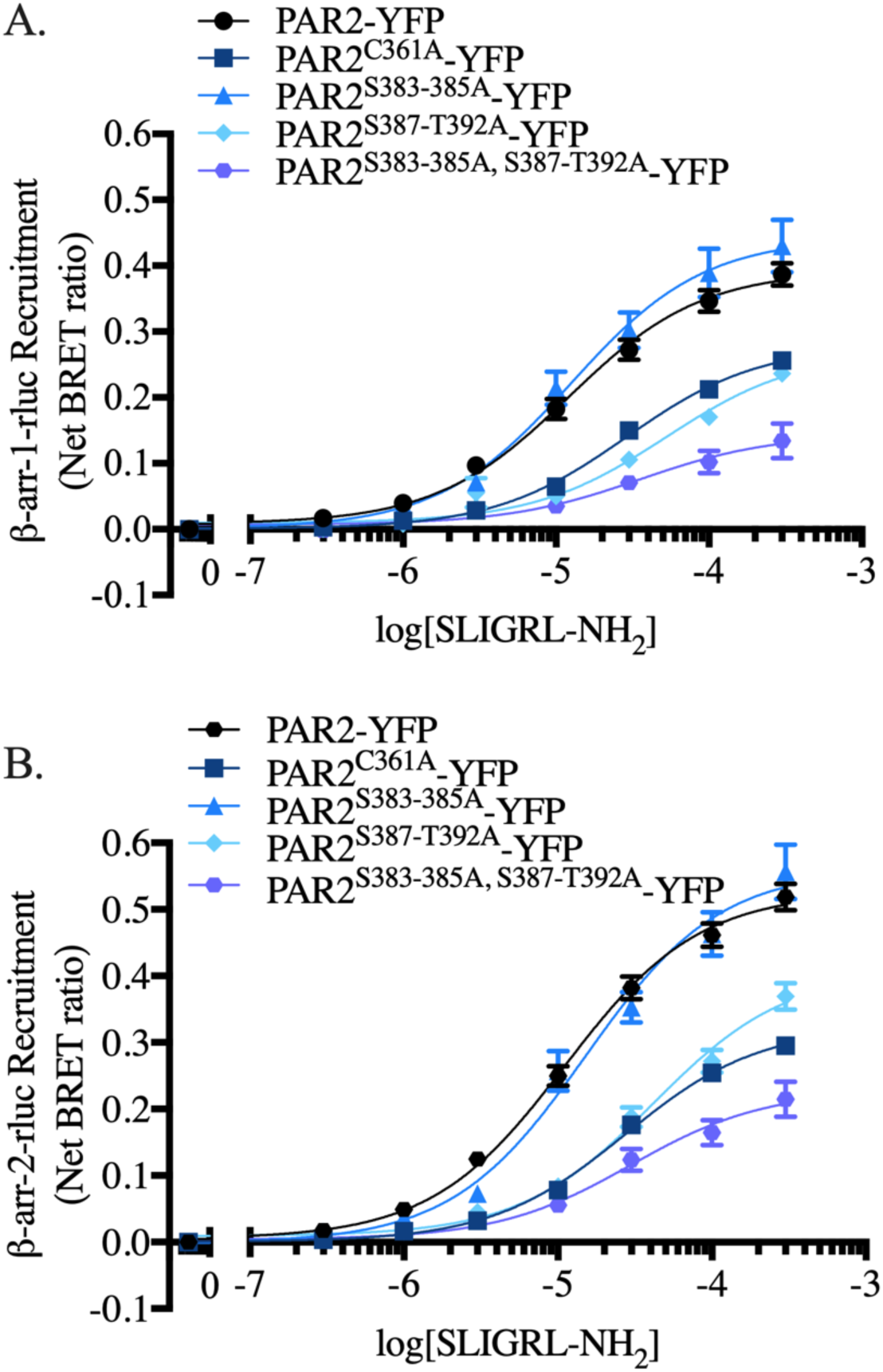
β-arrestin-1/-2 recruitment in PAR2 cysteine and phosphorylation mutants in response to SLIGRL-NH_2_ stimulation. β-arrestin recruitment concentration-effect curves for HEK-293 cells transiently expressing PAR2 or C-tail mutant PAR2 constructs (PAR2^C361A^-YFP, square; PAR2^S383-385A^-YFP, triangle; PAR2^S387-T392A^-YFP, diamond; PAR2^S383-385A,S387-T392A^-YFP, hexagon) with β-arrestin-1-rluc (A) or −2-rluc (B) were stimulated with PAR2 peptide agonist, SLIGRL-NH_2_. Nonlinear regression curve fits are shown (mean ± S.E.) for three to four independent experiments with triplicate data points for each concentration and receptor collected within each experiment. We find that Cys^361^Ala mutation significantly increased the concentration of SLIGRL-NH_2_ required to achieve half-maximal β-arrestin recruitment while concurrently demonstrating decreased maximal recruitment. Interestingly, Ser^383-385^Ala mutation did not significantly alter β-arrestin recruitment. Recruitment of β-arrestins was significantly decreased to both PAR2 Ser^387-^Thr^392^Ala and PAR2 Ser^383-385^Ala/Ser^387-^ Thr^392^Ala receptors. *(n = 3-4)*

Next, we evaluated mutations to phosphorylatable residues in the PAR2 C-tail. Mutation of Ser^383-385^ to alanine (PAR2^S383-385A^-YFP) did not significantly affect recruitment of β-arrestin-1 or −2 (EC_50_ = 12.5 ± 2.8 µM and 15.1 ± 2.7 µM, respectively). In addition, we observed no change in β-arrestin-2 recruitment Max. for PAR2^S383-385A^-YFP compared to wildtype PAR2. Maximum recruitment for β-arrestin-1 recruitment in this mutant was however significantly decreased (β-arrestin-1 Max. = 0.44 ± 0.02; β-arrestin-2 Max. = 0.56 ± 0.02). Mutation of Ser^387^-Thr^392^ to alanine (PAR2^S387-T392A^-YFP) significantly reduced both EC_50_ and Max. for both β-arrestin-1 (EC_50_ = 52.4 ± 12.1 µM; Max. = 0.27 ± 0.02) and β-arrestin-2 (EC_50_ = 41.7 ± 6.3 µM; Max. = 0.40 ± 0.02) compared to wild-type receptor. Interestingly, combined mutation of both Ser^383-385^ and Ser^387^-Thr^392^ to alanine (PAR2^S383-384A,S387-T392A^-YFP) was not significantly more detrimental to β-arrestin recruitment than Ser^387^-Thr^392^ to Ala alone. Recruitment of both β-arrestin-1 (EC_50_ = 35.4 ± 17.0 µM; Max. = 0.14 ± 0.02) and β-arrestin-2 (EC_50_ = 29.9 ± 8.6; Max. = 0.23 ± 0.02) were not significantly decreased compared to PAR2-YFP. (β-arrestin-1 BRET Fig. 2A; β-arrestin-2 BRET Fig. 2B)

### PAR2 β-arrestin recruitment is unaffected by C-tail phosphorylatable residue mutations in response to trypsin stimulation

To investigate the role of C-tail mutations on the recruitment of β-arrestins in response to tethered-ligand activation of PAR2, we employed BRET assays with PAR2 and PAR2 C-tail mutants stimulated with increasing concentrations of trypsin. Trypsin-cleavage reveals the PAR2 tethered ligand and thus allowed us to determine the impact of C-tail mutations on tethered ligand mediated PAR2 activation. First, we recorded agonist stimulated β-arrestin-1 and −2 recruitment with wildtype PAR2-YFP (Fig. 3). We observe a trypsin-stimulated, concentration-dependent, recruitment of β-arrestin-1 (EC_50_ = 9.4 ± 1.6 nM; Max. = 0.17 ± 0.01) and −2 (EC_50_ = 6.1 ± 1.2 nM; Max. = 0.14 ± 0.01). To investigate the known palmitoylation site of the PAR2 C-tail, we recorded β-arrestin recruitment to PAR2^C361A^-YFP following trypsin-stimulated activation of the receptor. We observed no significant difference in the potency and a modest decrease in the maximum for recruitment of β-arrestin 1 (EC_50_ = 5.7 ± 1.1 nM; Max. = 0.12 ± 0.01) and −2 (EC_50_ = 4.9 ± 0.7 nM; Max. = 0.12 ± 0.00) in this mutant.

**Figure 3:**
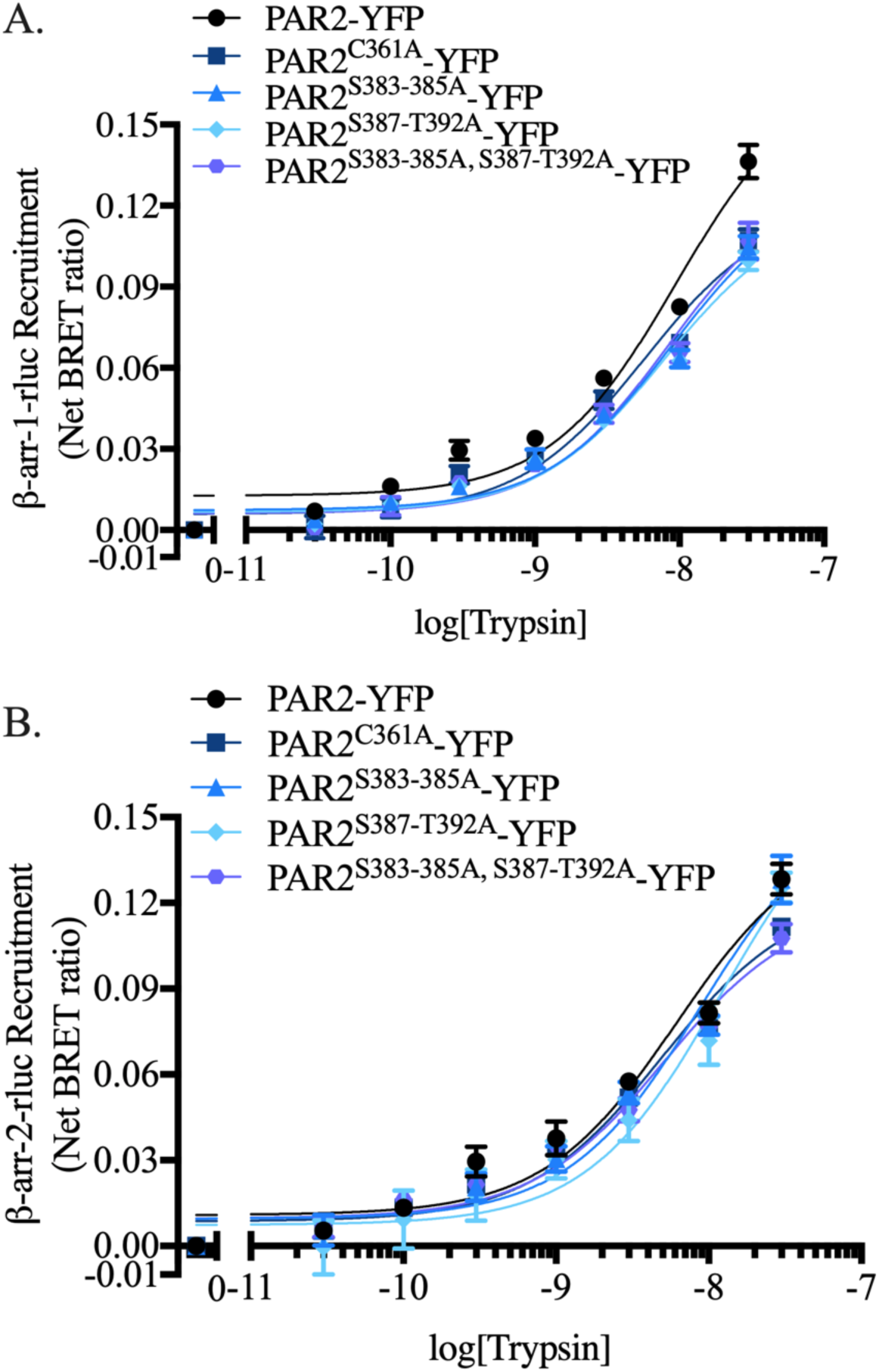
β-arrestin-1/-2 recruitment in PAR2 cysteine and phosphorylation mutants in response to tethered ligand agonism by trypsin stimulation. β-arrestin recruitment concentration-effect curves for HEK-293 cells transiently expressing PAR2 or C-tail mutant PAR2 constructs (PAR2^C361A^-YFP, square; PAR2^S383-385A^-YFP, triangle; PAR2^S387-T392A^-YFP, diamond; PAR2^S383-385A, S387-T392A^-YFP, hexagon) with β-arrestin-1-rluc (A) or −2-rluc (B) stimulated with trypsin. Nonlinear regression curve fits are shown (mean ± S.E.) for three to four independent experiments with triplicate data points for each concentration and receptor collected within each experiment. We find that no mutations resulted in significant shifts to EC_50_ of trypsin-stimulated β-arrestin-1 or −2 recruitment; however, significant decreases in maximal β-arrestin-1-rluc recruitment were observed with all mutants. *(n = 3-4)*

We then turned our investigation to the effect of mutations in C-tail phosphorylation sites and their impact on trypsin mediated β-arrestin recruitment. PAR2^S383-385A^-YFP was able to recruit β-arrestin-1 at half-maximal trypsin concentrations that were not significantly different than that observed with wildtype PAR2 (EC_50_ = 8.7 ± 1.6 nM). The maximum recruitment of β-arrestin-1 was however significantly reduced (Max. = 0.13 ± 0.01). β-arrestin-2 recruitment by trypsin activation of this mutant was comparable to that observed with wildtype PAR2 (EC_50_ = 8.8 ± 2.1; nM Max. = 0.16 ± 0.01). Similarly, in the case of PAR2^S387-T392A^-YFP stimulated with trypsin, β-arrestin recruitment occurred with EC_50_ values that were not significantly different than the wildtype receptor, however a significant decrease in Max. for β-arrestin-1 recruitment was also seen for this mutant (β-arrestin-1 EC_50_ = 7.8 ± 1.2 nM, Max. = 0.12 ± 0.01; β-arrestin-2 EC_50_ = 11.9 ± 4.6 nM, Max. = 0.17 ± 0.02). Additionally, we do not observe a significant alteration in β-arrestin recruitment compared to wildtype PAR2 with combined mutation of PAR2 Ser^383-385^ and Ser^387^-Thr^392^ to alanine (β-arrestin-1 EC_50_ = 8.4 ± 1.8 nM, Max. = 0.13 ± 0.01; β-arrestin-2 EC_50_ = 5.1 ± 1.0 nM, Max. = 0.12 ± 0.01).

In summary, mutations of the C-terminal Cys^361^ residue resulted in a decrease in β-arrestin recruitment to PAR2. Interestingly, this effect was more pronounced in the PAR2 activation with the synthetic peptide agonist compared to trypsin-mediated, tethered-ligand activation of the receptor. Mutations to clusters of serine and threonine residues show a clear role for the Ser^387^-Thr^392^ cluster, but not for the Ser^383-385^ cluster which was once again seen in the peptide-activated receptor but not with trypsin stimulation. (β-arrestin-1 BRET Fig. 3A; β-arrestin-2 BRET Fig. 3B)

### Agonist-stimulated β-arrestin recruitment can occur independent of Gα_q/11_ signalling

To determine whether Gα_q/11_ activation is necessary to stimulate recruitment of β-arrestins to PAR2, we monitored β-arrestin recruitment (BRET) following inhibition of Gα_q/11_ using YM254890 (Wako Chemicals). We observed a significant decrease in the maximal β-arrestin recruitment in cells pretreated with Gα_q/11_ inhibitor (YM254890, 100nM) compared to vehicle control (0.1% DMSO) following activation of PAR2 with SLIGRL-NH_2_ (β-arrestin-1 BRET Fig. 4A, β-arrestin-2 BRET Fig. 4B; statistical analysis Fig. 4E). In contrast, no significant changes in EC_50_ were observed (Fig. 4E). SLIGRL-NH_2_-stimulated β-arrestin-1 recruitment was determined to have an EC_50_ of 15.7 ± 3.6 µM in vehicle-treated controls and 20.9 ± 2.4 µM in YM254890 treated cells. β-arrestin-2 recruitment was determined to have an EC_50_ of 19.0 ± 2.8 µM in cells pre-incubated with vehicle control and 26.5 ± 3.2 µM in cells pretreated with YM254890.

**Figure 4:**
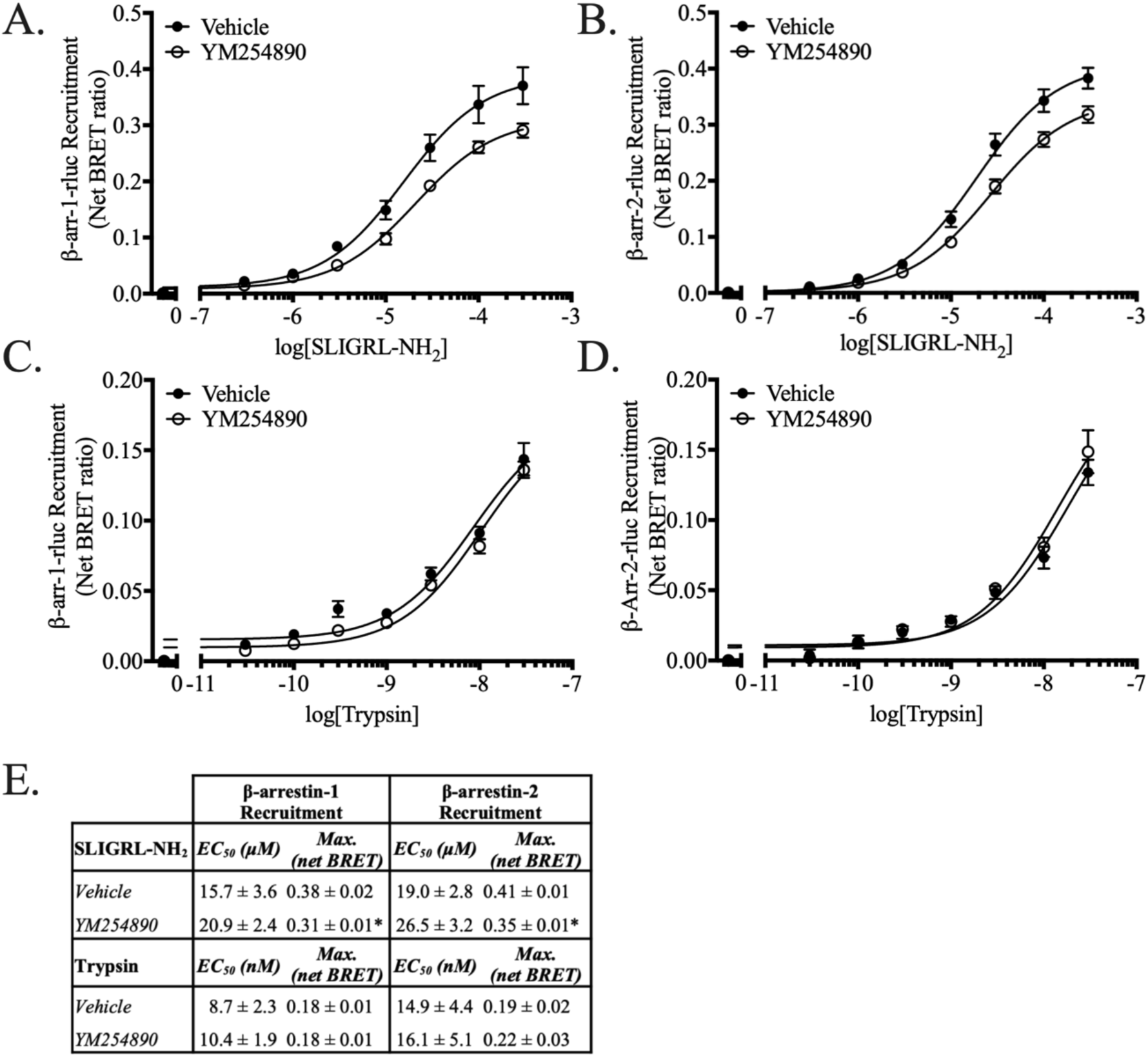
PAR2 agonist-dependent β-arrestin recruitment occurs independent of Gα_q/11_ activation. HEK-293 cells transiently expressing PAR2-YFP and either β-arrestin-1-rluc or −2-rluc were stimulated with increasing concentrations of either SLIGRL-NH_2_ (β-arr-1- (A), β-arr-2-rluc (B)) or trypsin (β-arr-1- (C), β-arr-2-rluc (D)) following preincubation with either vehicle control (DMSO, 0.01%; solid circles) or Gα_q/11_ selective inhibitor, YM254890 (100 nM; open circles). There were no significant shifts in EC_50_ between vehicle- or YM254890-treated following stimulation with either SLIGRL-NH_2_ or trypsin. Maximal recruitment of β-arrestin up (Max.) was significantly decreased in response to SLIGRL-NH_2_ but not trypsin stimulation. Nonlinear regression curve fits are shown for data collected in triplicates for each of at least three independent experiments (mean ± S.E.) for three to five independent experiments with triplicate data points collected in each experiment. (*n = 3-5*)

In the case of trypsin activation of PAR2, no differences were observed in β-arrestin-1 or −2 recruitment (Fig. 4C & 4D. We determined EC_50_ of trypsin-stimulated β-arrestin-1 recruitment to be 8.7 ± 2.3 nM in cells pretreated with vehicle control compared with 10.4 ± 1.9 nM in cells pretreated with YM254890. Trypsin-stimulated recruitment of β-arrestin-2 was determined to be 14.9 ± 4.4 nM in control cells compared to 16.1 ± 5.1 nM in inhibitor-treated cells. (Fig. 4E) Overall, these data indicate that β-arrestin recruitment to PAR2 can occur independently of Gα_q/11_ activation, with peptide-mediated and tethered-ligand mediated receptor activation showing only modest differences in Gα_q/11_ recruitment.

### PAR2-mediated calcium signalling is prolonged in β-arrestin-knockout HEK-293 cells in response to both SLIGRL-NH_2_ and trypsin

In order to probe an effect of β-arrestins on Gα_q/11_-mediated signalling, we compared SLIGRL-NH_2_- and trypsin-stimulated PAR2 signalling in HEK-293 cells and CRISPR/Cas9 β-arrestin-knockout HEK-293 cells endogenously expressing PAR2. We observed a significant increase in SLIGRL-NH_2_-stimulated calcium signalling (as a % of response from 3 μM A23187) in β-arrestin-knockout HEK-293 cells (Fig. 5B) compared to HEK-293 cells (Fig. 5A) (EC_50_ = 14.1 ± 3.3 μM and EC_50_ = 55.0 ± 13.8 μM, respectively; p < 0.05). In addition to an evident leftward shift in the concentration effect curve for calcium signalling (Fig. 5D), we analyzed the area under the curve for the response over 80 seconds post-agonist addition. Area under the curve analysis revealed a concomitant prolongation of calcium signalling duration in β-arrestin-knockout HEK-293 cells (Max. = 4718 ± 4, arbitrary units) compared to HEK-293 cells (Max. = 2870 ± 7, arbitrary units) (Fig. 5C). A similar trend was observed in response to PAR2 activation with trypsin with the area under the curve increased in β-arrestin-knockout HEK-293 cells (Max. = 4677 ± 2) compared to wild-type HEK-293 cells (Max. = 2287 ± 3, arbitrary units) (Fig. 5G). Unlike signalling stimulated by SLIGRL-NH_2_, potency of trypsin-stimulated calcium signalling (as a % of response from 3 μM A23187) was not significantly different than wildtype HEK-293 cells (Fig. 5H) (β-arrestin-knockout HEK-293 EC_50_ = 4.7 ± 0.5 nM; HEK-293 EC_50_ = 6.6 ± 1.4 nM; Fig. 5F & 5E, respectively). Although no shift in potency was observed, it is notable that there was an increase in the maximal calcium signal achieved with trypsin in β-arrestin-knockout HEK-293 compared to wildtype HEK-293 (β-arrestin-knockout HEK-293 Max. = 84.1 ± 1.7 % of A23187; HEK-293 Max. = 67.8 ± 2.9 % of A23187; p > 0.05).

**Figure 5:**
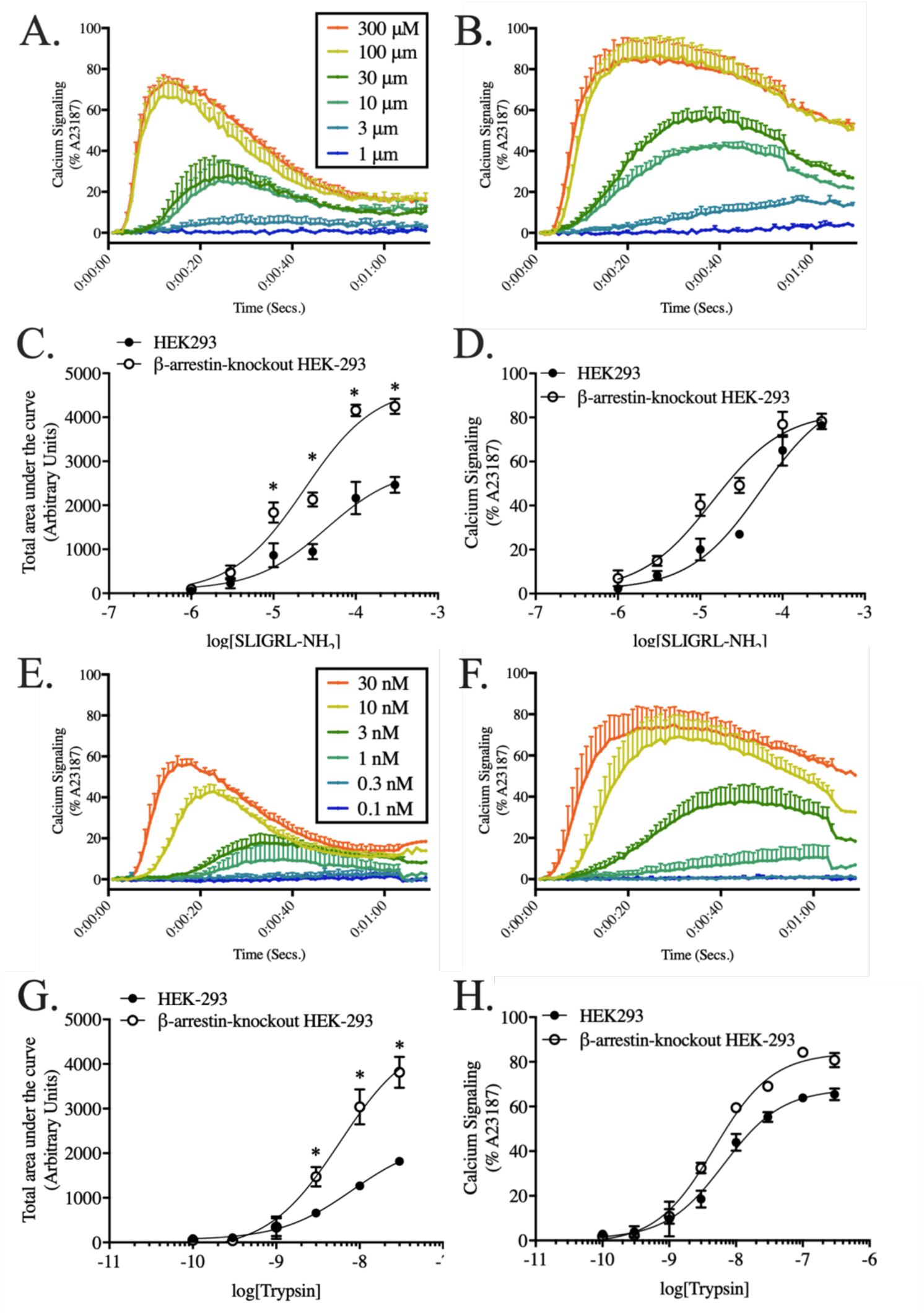
PAR2 agonist-dependent Gα_q/11_-mediated calcium signalling is prolonged and sensitized in β-arrestin-knockout HEK-293 cells compared to wildtype HEK-293 cells. SLIGRL-NH_2_-stimulated calcium signalling traces in HEK-293 (A) and β-arrestin-knockout HEK-293 (B) show prolonged signalling in β-arrestin-knockout HEK-293 cells. Both area under the curve analysis (C) and concentration-effect curve (D) of SLIGRL-NH_2_-stimulated calcium signalling reveal that signalling is prolonged and sensitized in β-arrestin-knockout HEK-293 cells compared to wildtype HEK-293 cells. Trypsin-stimulated calcium signalling traces in HEK-293 (E) and β-arrestin-knockout HEK-293 (F) show prolonged signalling in β-arrestin-knockout HEK-293 cells. Both area under the curve analysis (G) and concentration-effect curve (H) of trypsin-stimulated calcium signalling reveal that signalling is prolonged and sensitized in β-arrestin-knockout HEK-293 cells compared to wildtype HEK-293 cells. Calcium signalling is shown as a function of time, area under the curve, and maximal calcium signalling at given concentrations for three independent experiments. (*n = 3*)

### Differential calcium signalling in PAR2 C-tail mutants’ responses to SLIGRL-NH_2_ stimulation

To determine the importance of PAR2 C-tail palmitoylation and phosphorylation sites in calcium signal regulation, PAR2 C-tail mutants were expressed in PAR2-knockout HEK-293 cells (25) and calcium signalling was recorded in response to increasing concentrations of SLIGRL-NH_2_. Area under the curve was analyzed for recordings of 80 seconds post-agonist addition. For all receptor-mediated calcium signalling responses, both EC_50_ and maximum total area under the curve (“Max.”, 300 µM; arbitrary units) are reported with standard error. In response to agonist stimulation with SLIGRL-NH_2_ (0.3-300 µM), we observe responses with an EC_50_ = 1.1 ± 0.1 µM and Max. = 1433 ± 21 in the wild-type PAR2-YFP expressing cells (Fig. 6A & 6F). We observe a decreased SLIGRL-NH_2_ potency in the PAR2^C361A^-YFP expression (EC_50_ = 7.1 ± 0.9 µM) accompanied by a significantly increased maximal calcium signal (Max. = 1711 ± 37) (Fig. 6B & 6F).

**Figure 6:**
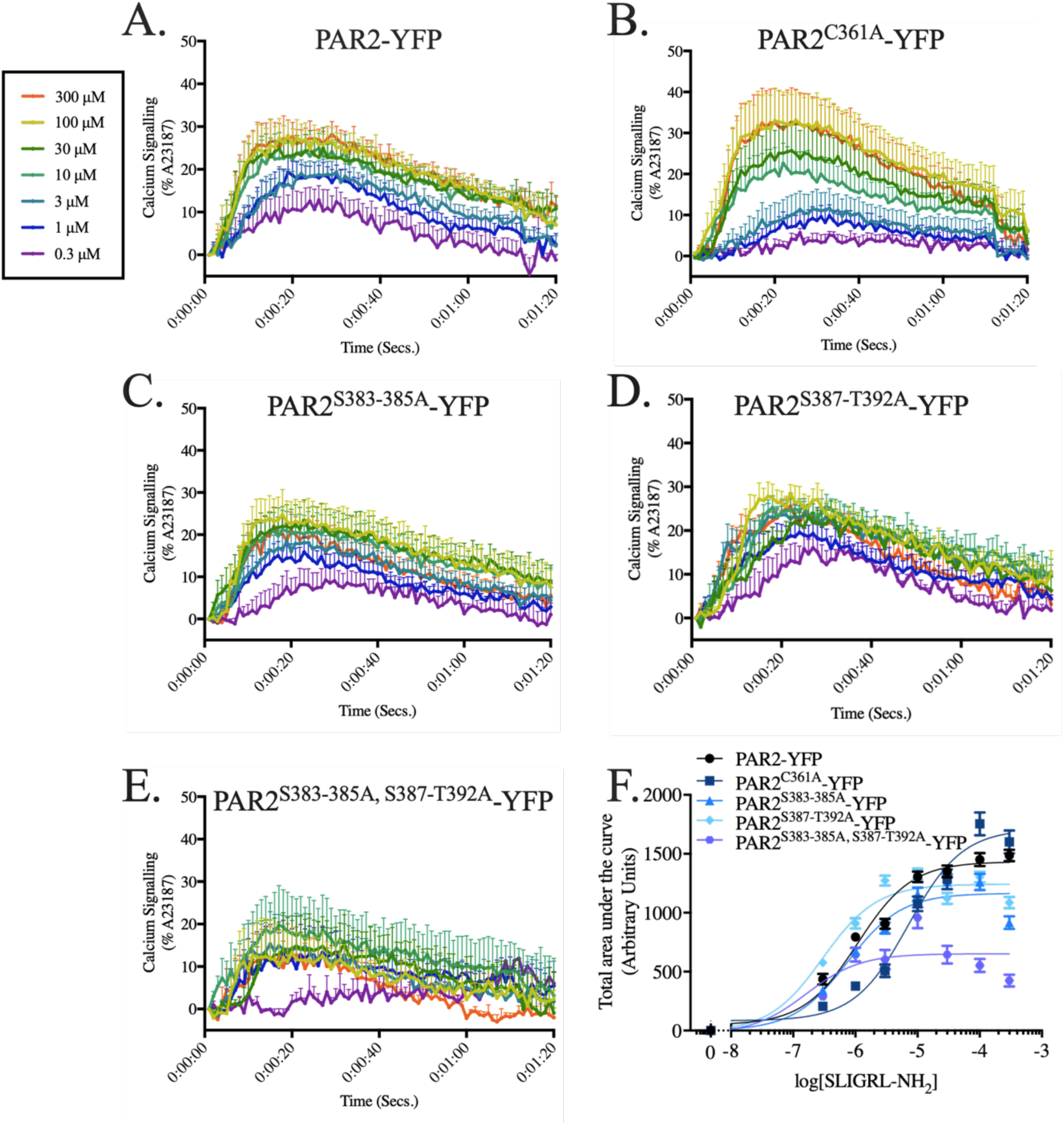
Differential calcium signalling in PAR2 mutant receptors compared to PAR2-YFP in response to stimulation with SLIGRL-NH_2_. Representative kinetic traces are shown for SLIGRL-NH_2_-stimulated intracellular calcium signalling (mean ± S.E.) for four to six independent experiments with PAR2-YFP (A), PAR2^C361A^-YFP (B), PAR2^S383-385A^-YFP (C), PAR2^S387-T392A^-YFP (D), and PAR2^S383-385A, S387-T392A^-YFP (E) receptors. Analysis of PAR2C^361^A-YFP mutant reveals significant dysregulation of calcium signalling compared to wild-type PAR2-YFP signalling. Serine mutations (Ser^387-^Thr^392^Ala and Ser^383-385^Ala/Ser^387-^Thr^392^Ala) modestly improve potency of SLIGRL-NH_2_ compared to PAR2-YFP. Interestingly, all serine/threonine mutants exhibited decreased maximal signalling elicited by SLIGRL-NH_2_ stimulation, with the most substantial decrease observed with the combined Ser./Thr. mutant (Ser^383-385^Ala/Ser^387-^Thr^392^Ala). Nonlinear regression curve fit of area under the curve shown for area under the curve (calculated with Prism 7) (F). (*n = 4-6*)

Calcium signalling in the C-tail phosphorylation site mutant, PAR2^S383-385A^-YFP, was no different than the wild-type receptor expressing cells (EC_50_ = 0.8 ± 0.1 µM) however there was a decrease in the maximal calcium signalling (Max. = 1165 ± 28) compared to PAR2-YFP (Fig. 6C & 6F). The other phosphorylation site mutant, PAR2^S387^-YFP, showed a modest but significant increase in calcium signalling (EC_50_ = 0.3 ± 0.1 µM) but also had decreased maximal calcium signal compared to the wild-type receptor (Max. = 1242 ± 23) (Fig. 6D & 6F). Combination of both Ser^383-385^Ala and Ser^387-T392^ Ala mutations (PAR2^S383-385A, S387-T392A^-YFP) had a cooperative effect, showing a greater increase in calcium signalling in response to SLIGRL-NH_2_ (EC_50_ = 0.2 ± 0.1 µM). This combined mutation, however, also resulted in a decrease in the total maximal signal elicited (Max. = 652 ± 44) (Fig. 6E & 6F). These results reveal that the loss of phosphorylatable residues sensitizes the calcium signal to SLIGRL-NH_2_ activation, suggesting that decreased regulation of receptor-mediated calcium signalling is responsible for the apparent leftward shift in potency observed in phosphorylation mutants. However, these C-terminal serine/threonine residues also appear to be important for calcium signalling since the deletion of both clusters significantly decreased the maximal calcium response following PAR2 activation.

### Alterations in calcium signalling with PAR2 C-tail mutants in response to trypsin agonism in PAR2 knockout HEK-293 cells

To investigate the importance of PAR2 C-tail palmitoylation site and phosphorylation sites in calcium signalling responses to receptor activation with the PAR2 tethered ligand, we examined calcium signalling following activation with trypsin. We recorded PAR2-YFP mediated calcium signalling in response to 0.03-30 nM trypsin and report an EC_50_ of 0.6 ± 0.2 nM and Max. of 1729 ± 125 (arbitrary units) (Fig. 7A & 7F). As observed with SLIGRL-NH_2_ stimulation, mutation of Cys^361^ to alanine resulted in a rightward shift in calcium response (EC_50_ = 2.3 ± 0.9 nM) and increased maximum total area compared to PAR2-YFP (Max. = 2081 ± 193) (Fig. 7B & 7C). This suggests that destabilization of the PAR2 C-tail through loss of a cysteine palmitoylation site causes dysregulation of calcium signalling resulting in less efficient activation of the pathway; while prolongation of signalling suggests less efficient desensitization of receptor-mediated calcium signalling.

**Figure 7:**
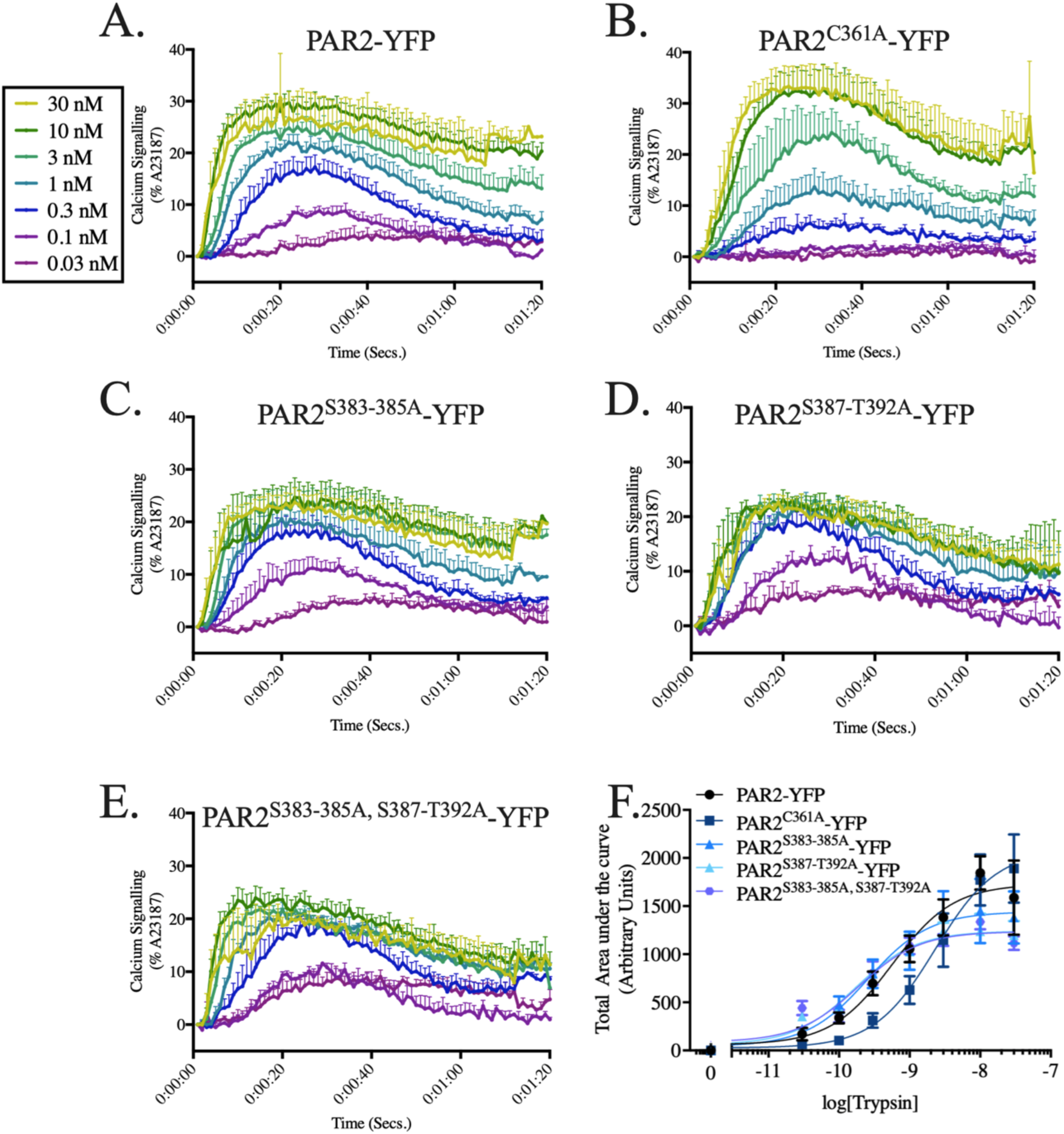
Differential calcium signalling in PAR2 mutant receptors compared to PAR2-YFP in response to stimulation with trypsin. Representative kinetic traces are shown for trypsin-stimulated intracellular calcium signalling (mean ± S.E.) for three independent experiments with PAR2-YFP (A), PAR2^C361A^-YFP (B), PAR2^S383-385A^-YFP (C), PAR2^S387-T392A^-YFP (D), and PAR2^S383-385A, S387-T392A^-YFP (E) receptors. Analysis of PAR2^C361A^-YFP mutant shows rightward shift in potency compared to wild-type PAR2-YFP signalling. As observed with calcium signalling in response to SLIGRL-NH_2_ stimulation, all PAR2 C-tail phosphorylation site mutant receptors were observed to have decreased maximal calcium signalling compared to PAR2-YFP. Nonlinear regression curve fit of area under the curve shown for area under the curve (calculated with Prism 7) (F). (*n = 3*)

To investigate the effect of phosphorylation site mutations on calcium signalling in response to tethered ligand activation following trypsin cleavage, we compared wild-type receptor and our three phosphorylation mutant receptors. In response to trypsin stimulation, we observe that PAR^S383-385A^-YFP results in both less potent activation of calcium signalling (EC_50_ = 2.6 ± 0.1 nM) as well as decreased maximum total area (Max. = 1442 ± 112) (Fig. 7C & 7F). Both PAR2^S387-T392A^-YFP (Fig. 7D) and combination mutant, PAR2^S383-385A,S387-T392A^-YFP (Fig. 7E) showed an increase in receptor-mediated calcium signalling in response to trypsin (EC_50_ = 0.2 ± 0.1 nM and EC_50_ = 0.2 ± 0.1 nM, respectively) but decreased the Max. compared to PAR2-YFP (Max. = 1241 ± 67 and Max. = 1236 ± 57, respectively) (Fig. 7F).

### Phosphorylation of ERK is stimulated by Gα_q/11_ and desensitized by β-arrestin

To investigate the contribution of PAR2 stimulated G-protein and β-arrestin signalling on the phosphorylation of ERK (p-44/42), we employed pharmacological inhibition of Gα_q/11_ with the compound YM254890 and used a CRISPR/Cas9 β-arrestin-knockout HEK-293 cell (26). Both pathways were explored as ERK phosphorylation can occur downstream of Gα_q/11_-dependent and/or β-arrestin-dependent/G-protein-independent pathways. Cells were stimulated with SLIGRL-NH_2_ (30 µM) for various time points to record ERK phosphorylation (p-p44/42). We observe that SLIGRL-NH_2_ mediated ERK phosphorylation was significantly inhibited in cells treated with 100 nM YM254890 prior to agonist stimulation (Vehicle AUC = 841 ± 183; YM254890 AUC = 206 ± 69) (Fig. 8A & 8C). Unexpectedly, we find that ERK signalling is greatly increased in β-arrestin double knockouts cells compared to HEK-293 cells (HEK-293 AUC = 812 ± 249; β-arrestin-knockout HEK-293 AUC = 2652 ± 855) (Fig. 8B & 8D). These data suggest that phosphorylation of ERK downstream of PAR2 stimulation of SLIGRL-NH_2_ is primarily mediated by Gα_q/11_ and subsequently desensitized/regulated by β-arrestin proteins.

**Figure 8:**
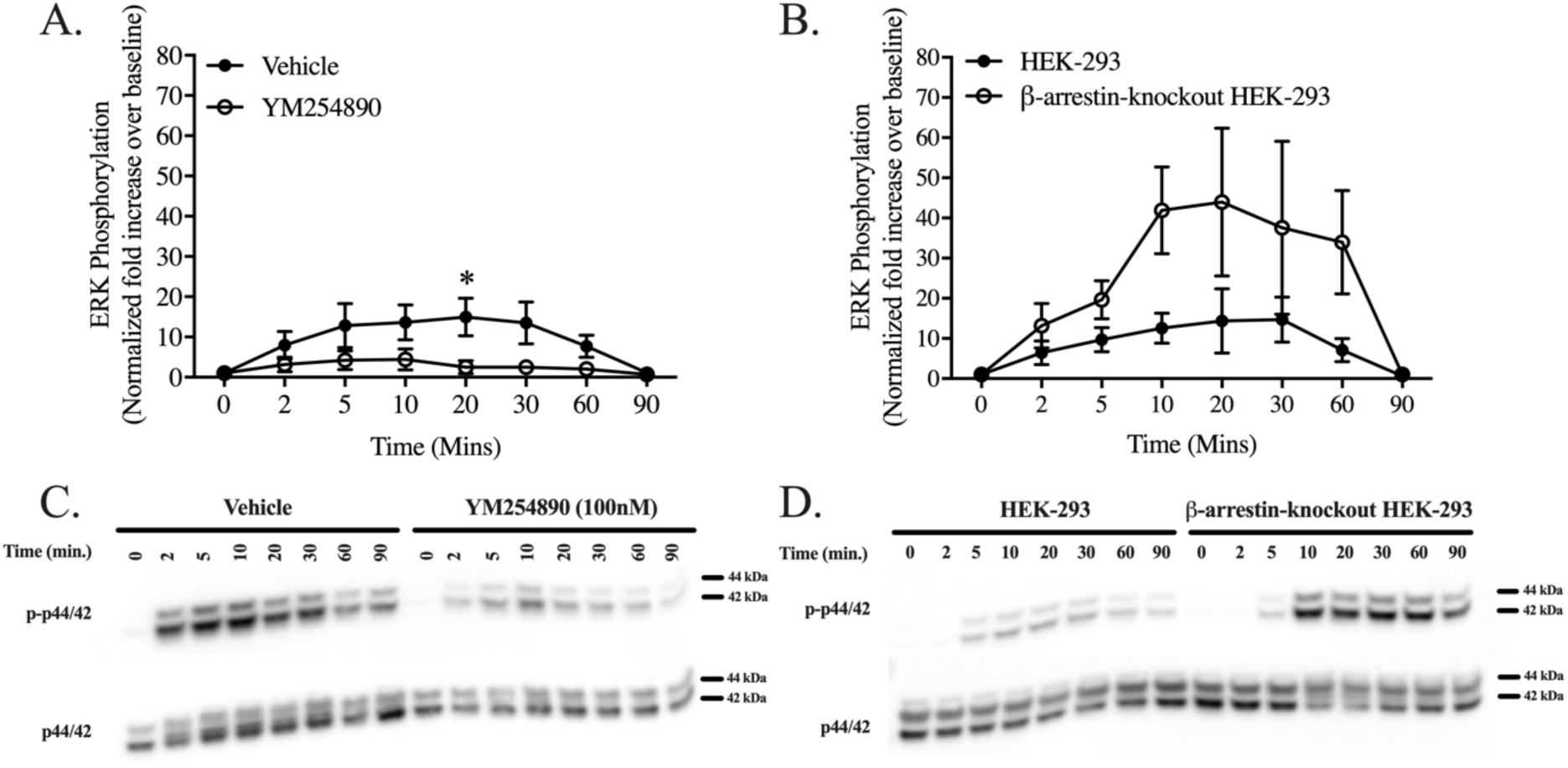
SLIGRL-NH_2_-stimulated, PAR2-mediated MAPK activation is Gα_q/11_-dependent and negatively regulated by β-arrestins. PAR2-mediated p44/42 phosphorylation is G**α**_**q/11**_ dependent. (A) SLIGRL-NH_2_-stimulated (30 µM) p44/42 phosphorylation over time in HEK-293 cells pre-treated with vehicle control (DMSO, 0.01%) or Gα_q/11_ inhibitor YM254890 (100 nM) for 20 minutes. (B) PAR2-mediated p44/42 phosphorylation in HEK-293 compared to β-arrestin-knockout HEK-293 cells following stimulation with SLIGRL-NH_2_ (30 µM). Analysis of p44/42 phosphorylation reveal that PAR2-stimualted MAPK activation is largely Gα_q/11_-dependent. Interestingly, normalized fold increase in p44/42 phosphorylation in HEK-293 and β-arrestin-knockout HEK-293 cells reveal that β-arrestin may be responsible for negative regulation/desensitization of PAR2-stimulated ERK phosphorylation. Data shown are representative of at least three independent experiments (phosphorylated to total p44/42, normalized to baseline phosphorylation at 0 minutes; mean ± S.E.; *p > 0.05) Representative blots of phosphorylated p44/42 (p-p44/42) and total p44/42 (p44/42) for vehicle/YM254890 (C) and HEK-293/β-arrestin-knockout HEK-293 (D) experiments are shown. (*n = 3-4*)

### Mutations of PAR2 C-tail stimulate increased levels of ERK phosphorylation

To investigate the effect of PAR2 C-tail mutations on phosphorylation of ERK we stimulated PAR2-knockout HEK-293 transiently expressing PAR2 C-tail mutants for 10 minutes with a sub-maximally activating concentration of SLIGRL-NH_2_ (3 μM) or trypsin (0.3nM). These concentrations were selected as they do not significantly activate ERK phosphorylation with wildtype receptor and therefore significant increases with mutant receptor would be apparent. Most importantly, since ERK signalling is downstream of receptor activation, it may be subject to amplification/deamplification processes that could mask significant increases in ERK phosphorylation downstream of mutant PAR2 receptor constructs at the highest concentrations used in this study. Interestingly, we observed an increase in ERK signalling [fold increase over baseline = (p-p44/42)/(p-44/42) normalized to untreated] with PAR2_C361A_-YFP (SLIGRL-NH_2_, 1.6 ± 0.3; trypsin, 1.2 ± 0.2), PAR2^S383-385A^-YFP (SLIGRL-NH_2_, 1.8 ± 0.4; trypsin, 1.3 ± 0.3), and PAR2^S387-T392A^-YFP (SLIGRL-NH_2_, 1.3 ±0.2; trypsin, 1.3 ± 0.3), compared to wildtype PAR2-YFP in response to SLIGRL-NH_2_ (3 µM; PAR2-YFP = 0.9 ± 0.2) and trypsin (0.3 nM; PAR2-YFP = 0.7 ± 0.1). Overall, we find that there is a significant increase in PAR2^S383-385A, S387-T392A^-YFP in response to both SLIGRL-NH_2_ (2.0 ± 0.3-fold increase over baseline) and trypsin (1.9 ± 0.6-fold increase over baseline) (Fig. 9A).

**Figure 9:**
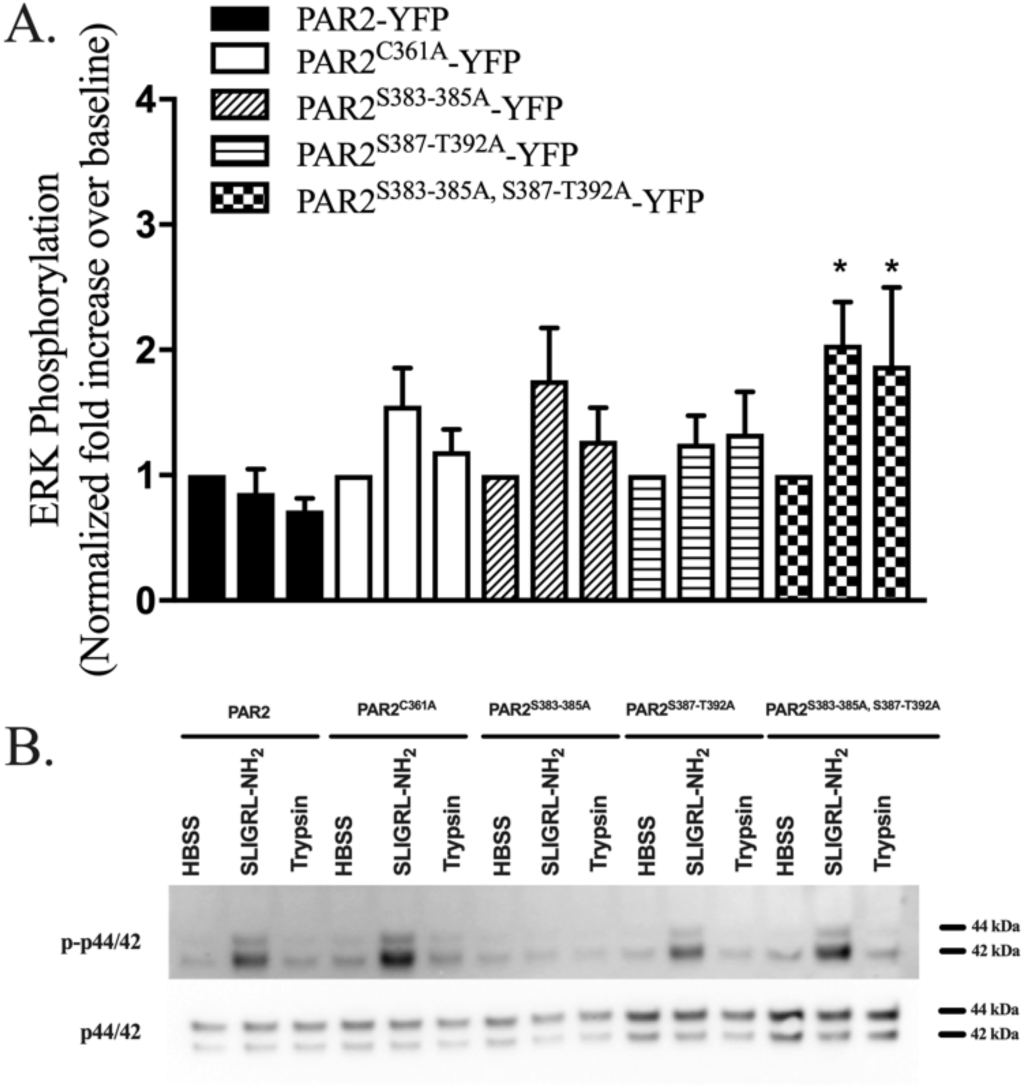
PAR2 C-tail phosphorylation mutation, Ser^383-385^Ala/Ser^385^-Thr^392^Ala, increases ERK phosphorylation in response to SLIGRL-NH_2_ and trypsin stimulation. (A) Summary data of p44/42 phosphorylation in PAR2-knockout HEK-293 cells expressing PAR2-YFP or PAR2 C-tail mutant receptors stimulated for 10 minutes with SLIGRL-NH_2_ (3 µM) or trypsin (0.3 nM). Sub-maximal concentrations of SLIGRL-NH_2_ and trypsin were chosen to avoid ERK activation with wildtype receptor to visual receptor mutations enhancing phosphorylation. Statistical analysis (two-way ANOVA) reveals that C-tail cysteine and phosphorylation mutants increase ERK phosphorylation compared to wildtype PAR2-YFP. Data shown are representative of four independent experiments (mean ± S.E.). (B) Representative blot of phosphorylated p44/42 (p-p44/42) and total p44/42 (p44/42) are shown. (*n = 4*)

## DISCUSSION

PAR2 is a proteolytically activated receptor that is a key regulator of inflammatory responses (27). The mechanisms that underlie biased signalling through PAR2 are of interest both for understanding receptor regulation of cellular behaviour and for developing therapeutic strategies that selectively target certain pathways. Here we examined determinants of PAR2 coupling to two key effectors, Gα_q/11_ and β-arrestins. Building on previous work that identified regulatory motifs in the PAR2 C-terminal tail that affect interaction with these effectors (21–23), we find that a helix-8 cysteine residue (Cys^361^) is involved in PAR2 interaction with both Gα_q/11_ and β-arrestins. Clusters of serine and threonine residues, that are sites of phosphorylation by GPCR kinases (GRKs), in contrast, regulated β-arrestin interaction with PAR2 but did not have a big effect on Gα_q/11_-coupled calcium signalling.

We examined whether β-arrestin-1 and −2 recruitment to PAR2 can occur independently of Gα_q/11_-dependent signalling pathways. β-arrestin recruitment was modestly reduced by pre-treatment with a Gα_q/11_-specific inhibitor in response to SLIGRL-NH_2_ stimulation, while there was no difference in β-arrestin recruitment to cells pretreated with vehicle or the Gα_q/11_ inhibitor when trypsin was the activating agonist. This suggests that PAR2 can recruit β-arrestins independently of Gα_q/11_, though coupling to other Gα proteins may still precede β-arrestin interactions with PAR2.

Conversely, we probed the role of β-arrestin-1/-2 in regulating PAR2 signalling to Gα_q/11_ by examining responses to endogenously expressed PAR2 in a β-arrestin-knockout HEK-293 cell background and compared these to responses observed in the wildtype HEK-293 background. We observed a significant increase in the Gα_q/11_-dependent calcium signal, with increases in both magnitude and duration of signal observed with both trypsin and SLIGRL-NH_2_. This prolongation of the signal duration is entirely in keeping with the well-established role of β-arrestins as regulators of GPCR desensitization (28, 29). Coupling of PAR2 to Gα_q/11_ is negatively-regulated/desensitized by β-arrestin-1/-2, therefore the leftward shift in the concentration effect curve for both PAR2 agonists studied here in the β-arrestin-knockout HEK-293 cells further corroborates a role for β-arrestins in desensitizing PAR2 (22, 30, 31).

In addition to G-protein-dependent pathways, a role for β-arrestins as a scaffold for p42/44 MAPK signalling has been demonstrated for a number of GPCRs (29), including PAR2 (11). We therefore examined this pathway with the new tools we have to probe Gα_q/11_ and β-arrestins. We find that Gα_q/11_-inhibitor, YM254890, significantly attenuated phosphorylation of PAR2-dependent p42/44 MAPK activation. Since YM254890 did not inhibit β-arrestin recruitment, this decrease in ERK activation appears to stem entirely from an inhibition of the Gα_q/11_ signalling pathway. Further, in agreement with our observation of enhanced Gα_q/11_ signalling in β-arrestin-knockout HEK-293 cells, we find that phosphorylation of p42/44 was significantly increased in β-arrestin-knockout HEK-293 cells compared to wildtype. These data demonstrate that in the cells studied here, Gα_q/11_ signalling through PAR2 leads to p42/44 MAPK activation, which is negatively regulated by β-arrestins.

Phosphorylation of the C-tail is known to increase β-arrestin affinity for GPCRs (18, 32, 33). Additionally, molecular and structural studies have established a role for the 8^th^ helix in enabling both visual arrestin and β-arrestin interactions with GPCRs (19, 34–36). Both of the motifs investigated in this study have been studied for PAR2 but in different cellular backgrounds (21–23). To probe the relative contributions of these two mechanisms, we examined G-protein and β-arrestin engagement to PAR2 mutants with serine/threonine to alanine substitution or a construct where the 8^th^ helix cysteine residue was mutated to alanine.

Mutation of Cys^361^ to alanine significantly decreased recruitment of β-arrestin-1/-2 following agonist stimulation with SLIGRL-NH_2_. The decrease in β-arrestin recruitment in the helix-8 cysteine mutant is in keeping with other GPCRs where this has been observed including the vasopressin receptor (37) and the thyrotrophin receptor (38). Intriguingly, β-arrestin recruitment to this mutant was largely unaffected when the receptor was activated with trypsin to reveal the tethered-ligand, exhibiting only a modestly reduced maximal recruitment of β-arrestins but no apparent shift in potency.

Since helix-8 is also reported to be a site for G-protein interactions with GPCRs, we examined Gα_q/11_-mediated calcium signalling to PAR2 in the helix-8 cysteine mutant. We observe that the Cys^361^Ala mutation results in significant dysregulation of calcium signalling. This is demonstrated by both a higher concentration of agonist required to elicit a half maximal response and also by prolonged signalling following stimulation. The decrease in potency of PAR2 activators may be indicative of decreased Gα_q/11_ interactions with this mutant, while the slow desensitization and elevated maxima may be coming from the deficiency in β-arrestin recruitment noted earlier. However, we do not see any difference in responses to SLIGRL-NH_2_ or trypsin as noted for maximal β-arrestin recruitment, aside from a decreased potency of activators, which leads us to favour the idea that the mutated receptor is coupling to Gα_q/11_ less efficiently. Such defects in G-protein-coupling have been noted for the β2-adrenergic receptor (39, 40), M1 muscarinic acetylcholine receptor (41), M2 muscarinic acetylcholine receptor (42), glucagon like peptide-1 receptor (43), 5-Hydroxytryptamine(1A) receptor (44), CCR2 receptors (45), and some odorant receptors (46) - to highlight a few examples. In the case of the M3 muscarinic acetylcholine receptor, chemical cross-linking studies have implicated helix-8 residues (8.49 and 8.52 according to Ballesteros-Weinstein GPCR numbering system) in making agonist-promoted contacts with the α4-β6 loop of Gα_q_ (47). In contrast, G-protein-mediated signalling through other GPCRs such as the α2A-adrenergic receptor (48) and the vasopressin V_1A_ receptor (49) is not affected by mutations to the C-terminal cysteine residues. In the case of the vasopressin receptor, receptor phosphorylation under both basal and agonist-stimulated conditions was abolished when the helix-8 cysteine residue was mutated (49), suggesting a role in recruitment of GPCR kinases (GRKs). Whether Cys^361^Ala in PAR2 similarly affects GRK-mediated phosphorylation to affect β-arrestin recruitment, is not yet known. Thus, the decrease in β-arrestin-recruitment to PAR2 in the helix-8 cysteine mutant could be occurring through a direct disruption of a β-arrestin contact site; or through a deficit in GRK interaction and consequent defect in phosphorylation of the activated receptor. A more direct role for helix-8 as a site for protein-protein interaction is likely for Gα_q/11_ and PAR2.

GRK-mediated phosphorylation of GPCRs is well established as a critical modification required for β-arrestin interactions (29, 50). Further, the pattern of phosphorylation at different serine and threonine residues is GRK specific and can stabilize different arrestin conformations at activated receptors (51). The C-tail in PAR2 contains multiple clusters of serine and threonine residues that are established as sites of phosphorylation (22). We focused on two clusters encompassing residues Ser^383-385^ and Ser^387^-Thr^392^. We also generated a construct with combined mutations of both these clusters to alanine. We find that the membrane proximal cluster of serine residues does not play a role in agonist dependent recruitment of β-arrestin nor does it regulate Gα_q/11_-mediated signalling through PAR2. In contrast, mutation of the second cluster of six serine and threonine residues (Ser^387^-Thr^392^) to alanine significantly decreased SLIGRL-NH_2_-stimulated β-arrestin recruitment. Interestingly, we once again noted a different effect when the receptor was activated with trypsin, with only a modest decrease in the maximal recruitment of β-arrestins seen. Calcium signalling through this mutant was increased compared to the wild-type receptor in response to both peptide and enzyme activation. Combined deletion of both clusters of phosphorylation sites phenocopied β-arrestin recruitment and calcium signalling patterns seen with mutations to Ser^387^-Thr^392^ alone. Overall, this suggests that activation of PAR2 results in phosphorylation of C-terminal six amino acid Ser/Thr residue cluster at residues 387-392. Only a ∼50% reduction in β-arrestin recruitment was seen with mutations studied here, indicating that additional phosphorylatable residues provide β-arrestin contacts (22). Further, the lack of effect on trypsin recruitment of β-arrestin to PAR2 seen here could be indicative of different GRKs being involved in phosphorylating PAR2 following activation by agonist peptides or the trypsin-revealed tethered-ligand.

β-arrestin recruitment to GPCRs is an important scaffold for MAP kinase signalling (29). A role for β-arrestin in regulating ERK signalling kinetics (52) and specific sub-cellular locations (11) is also reported. Here we observed an increase in ERK phosphorylation in a C-tail phosphorylation site mutant when activating with SLIGRL-NH2 and trypsin. Combined with the previously noted decrease in β-arrestin recruitment to these mutants, the increase in ERK signalling is compatible with a model where ERK activation occurs downstream of G-protein signalling and is negatively regulated by β-arrestins. However, the helix-8 cysteine mutant, which showed a deficit in Gα_q/11_ signalling and β-arrestin recruitment, also exhibits a modest increase in ERK signalling, suggesting that another effector may be involved in ERK signalling downstream of PAR2.

ERK activation by PAR2 is also the only response monitored here that showed an involvement for the membrane proximal serine cluster. Combined mutation of Ser^383-385^ and Ser^387^-Thr^392^ resulted in a greater increase in ERK activation than seen with mutations in each individual cluster. In this case, the increase in ERK activation is seen in the mutants which also showed reduction in β-arrestin recruitment when activated with SLIGRL-NH_2_ further supporting the ideas the ERK activation here is predominantly occurring through a G-protein-mediated pathway. Overall, mechanisms regulating ERK activation downstream of PAR2 are likely agonist and context specific as reported for other GPCRs (53, 54) and warrant further study.

Together these studies provide novel insights into regulation of effector-coupling at the C-terminal tail of PAR2 and highlight the role of helix-8 and clusters of phosphorylated serine and threonine residues. These findings further highlight the complexity of mechanisms regulating β-arrestin recruitment to GPCRs and, in the case of PAR2, point to differences in receptor activation by soluble peptide agonist and the tethered-ligand.

## Experimental Procedures

### Chemicals and Other Reagents

Porcine trypsin (catalogue no. T-7418; ∼14,900 units/mg) was purchased from Sigma-Aldrich (St. Louis, MI). A maximum specific activity of 20,000 units/mg was used to calculate the approximate molar concentration of porcine pancreatic trypsin in the incubation medium (1 unit/ml, ∼2 nM) as described previously. SLIGRL-NH_2_ (> 95% purity by HPLC/MS) was purchased from Genscript (Piscataway, NJ) or EZBiolabs (Carmel, IN). Agonists were prepared in 4-(2-hydroxyethyl)-1-piperazineethanesulfonic acid (HEPES, 25 mM). All chemicals were purchased from Millipore-Sigma, Thermo Fisher Scientific (Hampton, NH), or BioShop Canada, Inc. (Burlington, Ontario, Canada), unless otherwise stated.

### Molecular Cloning and Constructs

The plasmid encoding the human PAR2 receptor with an in frame enhanced yellow fluorescent protein fusion tag (PAR2-YFP) was constructed as previously described (55). The use of an in-frame, PAR2 C-terminal fusion of enhanced yellow fluorescent protein has been utilized in several previous studies which demonstrated that PAR2-YFP couples to its known signalling pathways appropriately (56–58). Plasmid DNA mutations in the C-terminus of PAR2 were created using QuikChange XL Multi Site-Directed Mutagenesis kit (Agilent Technologies, Mississauga, ON, Canada) to generate all mutants described in this study. All constructs were verified by sanger sequencing (London Regional Genomics Centre, University of Western Ontario).

### Cell Lines and Culture Conditions

All media and cell culture reagents were purchased from Thermo Fisher Scientific (Waltham, MA). Human embryonic kidney (HEK) cells (HEK-293; ATCC), PAR2-knockout HEK-293, and β-arrestin-1/-2-knockout HEK-293 (β-arrestin-knockout HEK-293) (26) cell lines were maintained in Dulbecco’s modified Eagle’s medium supplemented with 10% fetal bovine serum, 1% sodium pyruvate, and penicillin streptomycin solution (50,000 units penicillin, 50,000 μg streptomycin). Since trypsin activates the PARs, cells were routinely sub-cultured using enzyme-free isotonic phosphate-buffered saline (PBS) containing EDTA (1 mM). Cells were transfected with PAR2-YFP or mutated PAR2-YFP receptor vectors using calcium phosphate [Nuclease-free water, 2.5M CaCl_2_, and 2x HEPES-buffered saline (HBS) (59)] or Fugene6 transfection methods (Promega, Madison, WI). Transiently transfected cells were always assayed or imaged at 48 hours post-transfection.

### Generation of CRISPR/Cas9 PAR2-knockout HEK-293

HEK-293 cells constitutively express PAR2 (60). In order to study mutant receptors in a null-background we generated PAR2-knockout HEK-293 cells using CRISPR/Cas9 targeting. The knockout design and procedures used to derive PAR2-knockout HEK-293 cells from the wild-type HEK-293 cells were as previously described (61, 62). A PAR2 specific guide (CCCCAGCAGCCACGCCGCGC) was cloned into the lentiCRISPR v2 plasmid (Addgene plasmid # 52961). 48 hours after transfection, cells were selected in media containing puromycin (5 µg/mL). PAR2-deficient cells were identified by functional screening of responses to PAR2 specific agonists using a calcium signalling assay (Figure S1).

### Calcium Signalling

Agonist-stimulated calcium signalling was recorded in PAR2-knockout HEK-293 cells as previously described (26, 55, 63). Cells were detached in enzyme-free cell dissociation buffer, centrifuged to pellet (1000 rpm, 5 minutes), and resuspended in Fluo-4 NW (no wash) calcium indicator dye (Thermo Fisher Scientific). Following a 30-minute incubation at ambient temperature, intracellular fluorescence (excitation 480 nm; emission recorded at 530 nm) was monitored before and after addition of agonists (trypsin or SLIGRL-NH_2_) on a PTI spectrophotometer (Photon Technology International, Birmingham, NJ). Responses were normalized to the fluorescence obtained with calcium ionophore (A23187, 3 μM; Sigma-Aldrich).

### Bioluminescence Resonance Energy Transfer Detection of β-arrestin-1/2 Recruitment

Bioluminescence resonance energy transfer (BRET) based detection of β-arrestin-1/-2 interaction with PAR2-YFP and mutant PAR2-YFP constructs was monitored in HEK-293 cells as described (14, 55). PAR2-YFP or mutant PAR2-YFP constructs (1 µg) and Renilla luciferase-tagged β-arrestin-1 or −2 (β-arr-1 and −2-rluc; 0.1 µg) were transiently transfected for 48 hours. Cells were plated in white 96-well culture plates (Corning; Oneonta, NY) and interactions between receptor and β-arrestin-1/-2 were detected by measuring BRET following 20 minutes of agonist stimulation and the addition of 5 μM coelenterazine prior to BRET recording (26) (NanoLight Technology, Pinetop, AZ) on a Mithras LB940 plate reader (Berthold Technologies, Bad Wildbad, Germany).

### Mitogen-Activated Protein Kinase Western BlotAssay

Agonist-stimulated mitogen-activated protein kinase (MAPK) signalling in PAR2-knockout HEK-293 cells expressing PAR2-YFP or mutant PAR2-YFP was monitored by Western blot analysis as previously described (26, 55, 63). Cells expressing PAR2 receptor constructs were serum starved (2 hours) and stimulated with sub-maximal concentrations of SLIGRL-NH_2_ (3 µM) or trypsin (0.3 nM) for 10 minutes at 37°C. Protein lysates were separated on 4-12% Bis-Tris gels (Invitrogen, Thermo Fisher Scientific). Phosphorylated p44/42 (p-p44/42; activated ERK) and total p44/42 (p44/42; total ERK) proteins were detected with antibodies (9101S and 9102S antibody clones, respectively, Cell Signalling Technology, Danvers, MA; 1:1000 concentration) and imaged by use of HRP-conjugated secondary antibodies (anti-rabbit IgG HRP-linked 7074S clone; Cell Signalling Technology, Danvers, MA; 1:10,000 concentration). Chemiluminescence was recorded on an iBright CL1000 gel doc (Invitrogen, Thermo Fisher Scientific) following application of ECL Prime detection reagent (GE Healthcare). Band intensities representing activated and inactive MAPK ERK proteins were quantified using the FIJI is just ImageJ (FIJI) quantification software (64). Phospho-kinase levels were normalized by expressing the data as a percentage of the corresponding total-kinase signal. Fold increase above baseline was calculated by normalizing data to baseline p44/42 phosphorylation in unstimulated samples.

### Assessing the contribution of Gα_q/11_ and β-arrestin-1/2 in MAPK activation

To examine the contribution of Gα_q/11_ signalling on PAR2-dependent MAPK activation, HEK-293, endogenously expressing PAR2 were pretreated with either vehicle control (DMSO, 0.01%) or YM254890 (100 nM) (65) for 20 minutes prior to agonist stimulation with SLIGRL-NH_2_ (30 µM). To assess the contribution of β-arrestin-1/-2 signalling in PAR2-dependent MAPK activation, HEK-293 and β-arrestin-knockout HEK-293 cells (26) endogenously expressing PAR2 were assayed for p44/42 phosphorylation following stimulation with SLIGRL-NH_2_ (30 µM).

### Confocal Microscopy

HEK-293 cells transiently transfected with PAR2-YFP or mutant PAR2-YFP (Fugene 6, Promega) were sub-cultured onto 35-mm glass-bottom culture dishes (MatTek Corporation, Ashland, MA) to be analyzed by confocal microscopy. Cells were fixed with 4% w/v paraformaldehyde solution, stained with 4′,6-diamidino-2-phenylindole (DAPI) to identify the nucleus, and receptor localization to the plasma membrane was assessed by imaging eYFP expression with an Olympus FV1000 (Centre Valley, PA) or Leica SP8 confocal microscope system (Buffalo Grove, IL) (Figure S2).

### Statistical Analysis

Statistical analysis of data, curve fitting (three-parameter nonlinear regression), and area under the curve analyses were done with Prism 7 software (GraphPad Software, La Jolla, CA). Statistical significance of EC_50_ shifts and concentration-effect curve top (indicated as “Max.”) were calculated using the extra sum of squares analysis and indicated in table 1 (*p < 0.05; Table 1) (26, 66). Statistical significance for western blots was assessed using two-way analysis of variance (*p < 0.05). Data are expressed as mean ± S.E. throughout the text, table, and figure legends.

**Table 1.** Summary of data β-arrestin-recruitment, calcium signalling, and MAPK activation with PAR2-YFP and PAR2-YFP mutants with SLIGRL-NH_2_ (top) and trypsin (bottom) stimulation. β-arrestin recruitment (EC_50_; “Max.” denotes maximum BRET signal obtained with a mutant up 300 µM SLIGRL-NH_2_ or 300 nM trypsin), calcium signalling [EC_50_; “Max.” denotes maximum calcium signal at 300 µM SLIGRL-NH_2_ or 300 nM trypsin (area under the curve; arbitrary units)], and phosphorylation of ERK (phosphorylated p44/42 / total p44/42; expressed as fold increase over unstimulated baseline) for PAR2-YFP and C-tail mutant PAR2-YFP receptors in response to SLIGRL-NH_2_ and trypsin. Statistical significance (* p < 0.05) was determined by least sum of squares analysis (EC_50_ and Max.) or two-way ANOVA (ERK phosphorylation).

|  | ***β*-arrestin-1**<br>**Recruitment** |  | ***β*-arrestin-2**<br>**Recruitment** |  | **Calcium** |  | **ERK Phosphorylation** |
| --- | --- | --- | --- | --- | --- | --- | --- |
| **SLIGRL-NH<sub>2</sub>** | ***EC*<sub>50</sub> (μM)** | **Max.**<br>**(net BRET)** | ***EC*<sub>50</sub> (μM)** | **Max.**<br>**(net BRET)** | ***EC*<sub>50</sub> (μM)** | **Max.**<br>**(A.U.C.)** | **Normalized fold increase**<br>**over baseline** |
| PAR2-YFP | 12.1 ± 1.6 | 0.39 ± 0.01 | 11.1 ± 1.2 | 0.53 ± 0.01 | 1.1 ± 0.1 | 1433 ± 21 | 0.9 ± 0.2 |
| PAR2<sup>C361A</sup>-YFP | 28.8 ± 4.0* | 0.28 ± 0.01* | 27.4 ± 2.5* | 0.32 ± 0.01* | 7.1 ± 0.9* | 1711 ± 37* | 1.6 ± 0.3 |
| PAR2<sup>S383-385A</sup>-YFP | 12.5 ± 2.8 | 0.44 ± 0.02* | 15.1 ± 2.7 | 0.56 ± 0.02 | 0.8 ± 0.1 | 1165 ± 28* | 1.8 ± 0.4 |
| PAR2<sup>S387-T392A</sup>-YFP | 52.4 ± 12.1* | 0.27 ± 0.02* | 41.7 ± 6.3* | 0.40 ± 0.02* | 0.3 ± 0.1* | 1242 ± 23* | 1.3 ± 0.2 |
| PAR2<sup>S383-385A, S387-T392A</sup>-YFP | 35.4 ± 17.0 | 0.14 ± 0.02 | 29.9 ± 8.6 | 0.23 ± 0.02 | 0.2 ± 0.1* | 652 ± 44* | 2.0 ± 0.3* |
| **Trypsin** | ***EC*<sub>50</sub> (nM)** | **Max.**<br>**(net BRET)** | ***EC*<sub>50</sub> (nM)** | **Max.**<br>**(net BRET)** | ***EC*<sub>50</sub> (nM)** | **Max.**<br>**(A.U.C.)** | **Normalized fold increase**<br>**over baseline** |
| PAR2-YFP | 9.4 ± 1.6 | 0.17 ± 0.01 | 6.1 ± 1.2 | 0.14 ± 0.01 | 0.6 ± 0.2 | 1729 ± 125 | 0.7 ± 0.1 |
| PAR2<sup>C361A</sup>-YFP | 5.7 ± 1.1 | 0.12 ± 0.01* | 4.9 ± 0.7 | 0.12 ± 0.00 | 2.3 ± 0.9* | 2081 ± 192 | 1.2 ± 0.2 |
| PAR2<sup>S383-385A</sup>-YFP | 8.7 ± 1.6 | 0.13 ± 0.01* | 8.8 ± 2.1 | 0.16 ± 0.01 | 2.6 ± 0.1 | 1442 ± 112 | 1.3 ± 0.3 |
| PAR2<sup>S387-T392A</sup>-YFP | 7.8 ± 1.2 | 0.12 ± 0.01* | 11.9 ± 4.6 | 0.17 ± 0.02 | 0.2 ± 0.1 | 1241 ± 67* | 1.3 ± 0.3 |
| PAR2<sup>S383-385A, S387-T392A</sup>-YFP | 8.4 ± 1.8 | 0.13 ± 0.01* | 5.1 ± 1.0 | 0.12 ± 0.01 | 0.2 ± 0.1 | 1236 ± 57* | 1.9 ± 0.6* |

## Acknowledgments

We would like to thank Dr. Michel Bouvier (U de Montreal) for providing the Renilla luciferase-tagged β-arrestin-1/-2 constructs and Dr. Feng Zhang (MIT) for providing the lentiCRISPR-V2 plasmid.

## FOOTNOTES

These studies were funded by a Canadian Institutes of Health Research (CIHR # 376560) grant (RR). PT is the recipient of a doctoral QEII Graduate Scholarship in Science and Technology.

## The abbreviations used are

PAR2: proteinase activated receptor-2
GPCR: G-protein-coupled receptor
MAPK: mitogen-activated protein kinase
ERK: extracellular-regulated protein kinase (p44/42)

